# Beyond abstract selection coefficients: Protandry impacts the buildup of heterozygote advantage over the lifespan in a color polymorphic moth

**DOI:** 10.64898/2026.03.29.715091

**Authors:** Eetu Selenius, Thomas Keaney, Sandra Winters, Johanna Mappes, Hanna Kokko

**Affiliations:** Organismal and Evolutionary Biology Research Programme, Faculty of Biological and Environmental Sciences, University of Helsinki, Viikinkaari 1, 00790, Helsinki Finland; Institute of Organismic and Molecular Evolution (iomE), Johannes Gutenberg University of Mainz, Mainz, Germany; Institute for Quantitative and Computational Biosciences (IQCB), Johannes Gutenberg University of Mainz, Mainz, Germany; BioFrontiers Institute, University of Colorado Boulder, Boulder, Colorado, USA

## Abstract

Population genetic models excel at identifying the conditions for polymorphisms based on balancing selection but typically disregard the ecological processes that yield particular values of selection coefficients. We model a system that combines antagonistic pleiotropy, dominance reversal and heterozygote advantage: the wood tiger moth *Arctia plantaginis*, where alternative haplotypes at a major-effect locus determine male hindwing coloration. Yellow offers better protection against predators, while white is often associated with better mating success. The effects of mortality and reproductive success overlap in time because protandrous males can mate as long as they are alive, but they need to avoid predation for several days before the bulk of females emerge. We show that protandry aids polymorphism maintenance whenever the second-fittest genotype (after the heterozygote) is the poorly surviving but mating advantaged homozygote, while increased protandry harms polymorphism when the second-best fitness is that of the survival advantaged morph. Ecologically plausible protandry times predict that dominance reversal does not have to be strong for polymorphism to be maintained. Our study highlights the importance of timing traits in maintaining polymorphisms in *Lepidoptera* and showcases the benefits of deriving fitness explicitly in place of abstract selection coefficients that lack temporal components within the life cycle.

## Introduction

Genetic polymorphisms and their maintenance have been topics of interest for evolutionary biologists for more than a century (Ruzicka et al. 2025). Some genetic variation can be explained by a balance between the removal of variation by selection and an input through mutation (Kimura and Crow 1964; Muller 1950), yet this is an insufficient explanation for the magnitude of fitness-affecting standing variation commonly observed in natural populations (Bonnet et al. 2022; Hendry et al. 2018; Keaney and Holman 2025). This abundance of genetic variance suggests the presence of additional maintenance mechanisms, of which balancing selection — where selection holds allele frequencies at an intermediate equilibrium — is the most popular (Hedrick et al. 1976; Ruzicka et al. 2025). A variety of processes have been found to generate balancing selection, including alleles having strong beneficial fitness effects in one environment, but negative ones in another (where environments can be niches, seasons, sexes, etc.; Bergland et al. 2014; Hedrick et al. 1976; Hoekstra et al. 2004; Kaufmann et al. 2023), rare genotypes being preferred in mate-choice contexts (Potter et al. 2023; Hedrick et al. 2016), and heterozygotes having higher fitness than homozygotes (Gemmell and Slate 2006). The latter effect can arise when a gene has antagonistically pleiotropic effects on traits expressed within a single organism, and the heterozygote expresses the favorable allele for both traits simultaneously (Curtsinger et al. 1994; Hedrick 1999), a phenomenon referred to as a beneficial reversal of dominance (Connallon and Chenoweth 2019; Rose 1982).

Theoretical models of polymorphism maintenance by antagonistic pleiotropy indicate that either large and approximately equal fitness effects or beneficial reversal of dominance are required to produce heterozygote advantage (Curtsinger et al. 1994; Gillespie 1976; Grieshop et al. 2024; Hedrick 1999). Historically, both conditions were considered unrealistic: large fitness effects would require near lethality and sterility, and both metabolic models and empirical data suggested dominance reversal to be unlikely in nature (Keightley and Kacser 1987; Curtsinger et al. 1994). Recently, insect laboratory studies indicate that dominance reversal can be wide-spread throughout the genome (Grieshop and Arnqvist 2018) and that it is associated with polymorphism in several species (Jardine et al. 2021; Mérot et al. 2020). This evidence, paired with dramatic examples of polymorphism maintenance from wild vertebrate populations (Pearse et al. 2019; Johnston et al. 2013; Barson et al. 2015), demonstrates that beneficial reversals of dominance can and do occur at fitness-affecting loci, overturning historical skepticism of their plausibility (Grieshop et al. 2024; Connallon and Chenoweth 2019).

Much of theory on balancing selection deals with abstract selection coefficients without deriving them from an explicit ecological setting (Hedrick 1999; Curtsinger et al. 1994; Rose 1982). While these approaches are useful in part due to their simplicity, we aim to show that valuable insights can be gained when the ecological underpinning is explicitly modelled. We explore how species-specific ecology (see also Brown and Kelly 2018; Tellier et al. 2007) can yield dominance reversal and heterozygote advantage in a situation where the life cycle of an organism does not permit a clean partitioning of selection coefficients that relate to survival separately from reproduction or vice versa.

The wood tiger moth (*Arctia plantaginis*), a model organism for polymorphic warning coloration, provides an interesting study system for antagonistic pleiotropy: it combines antagonistic pleiotropy in males with the potential for heterozygote advantage (De Pasqual et al. 2022). Wood tiger moths have male-limited polymorphic hindwing coloration, which is controlled by a large-effect autosomal locus in the yellow gene cluster containing the duplicated gene *valkea*, with two alternative haplotypes corresponding to the white (W) and yellow (y) morphs (Brien et al. 2023; Nokelainen et al. 2022). Males with one or two copies of the W allele appear white to the human eye but are distinctive to visual systems with ultraviolet sensitivity including the moths and their avian predators (Nokelainen et al. 2022) – suggesting that the W allele is not completely dominant. Males carrying two y alleles have yellow hind wing coloration (fig. S1).

Wing coloration impacts at least two adult fitness components: predator avoidance and mating, with a fitness trade-off between them. Yellow wings provide more effective warning coloration against avian predators and in some experiments are associated with stronger chemical defense, whereas white wings are linked to higher mating success (Nokelainen et al. 2012, 2014; Selenius et al. 2025; Rojas et al. 2017). Heterozygotes may therefore produce a ‘best of both worlds’ phenotype via dominance reversal if their version of white is at least somewhat aposematic to predators, while not suffering fully the cost of reduced mating success that would come with yellow wings, as suggested by previous research (Selenius et al. 2025). Critically, predation on adult male wood tiger moths is obviously deleterious to fitness, but it does not remove reproductive success already accrued. Here, we build models that treat both predation and matings as events that can occur throughout a male’s adult life, without enforcing any predetermined order to which these events must occur. This framework allows us to study the effect of protandry, where males enter the mating pool earlier on average than females, on polymorphism maintenance.

Protandry is a common feature of avian and insect mating systems (Wiklund and Fagerström 1977), including Lepidopterans with known sex-limited color polymorphisms such as members the *Colias* genus (Wiklund and Forsberg 1991; Klein et al. 2025). For insects, its evolutionary stability results from asymmetric fitness consequences when shifting emergence to earlier or later. Early-emerging males gain at least probabilistic access to females across the entire emergence period of females while late males definitely have missed all mating opportunities earlier than their own emergence (Ekrem and Kokko 2023; Zonneveld and Metz 1991). However, early emerging males are also exposed to prolonged periods of predation before the bulk of mating opportunities arise and therefore face significant risk of death before reproduction is possible. Therefore, it appears important to consider protandry as a factor that can intensify selection on survival-related fitness components (in wood tiger moths, the protective function of coloration). We show that this translates to either a broader or narrower region of dominance coefficients that permit a polymorphism, depending on the relative fitness ranking of the two homozygotes.

We first explore the effects of antagonistic pleiotropy, dominance reversals, and protandry generally with a stylized population genetic model. This version of the model is inspired by *A. plantaginis* but does not attempt a direct link to field data and oversimplifies protandry by assuming that all individuals of a given sex emerge simultaneously. The justification for these simplifications is the insight that can be gained through closed-form analytical solutions. We then switch to individual-based simulations with a more realistic emergence time distribution to investigate the maintenance of variation at color-affecting loci in *A. plantaginis*, parameterized with data from field experiments (Nokelainen et al. 2012; Selenius et al. 2025).

### Population genetic model

We first built a simple population genetic model to assess how protandry and antagonistic pleiotropy interact and use it to ask when these forces produce balancing selection at autosomal loci. We consider a panmictic, infinitely sized population with discrete generations and an even sex-ratio. Our focus is a single diploid locus segregating for two alleles: one which lowers the adult predation rate but decreases male mating success, and a second that produces the reverse effect. To match the fitness effects observed at the *valkea* gene in *A. plantaginis*, we refer to these as the y and W alleles, respectively. We further assume that expression of both y and W is limited to adult males and that the locus segregates in Mendelian fashion. Within a generation, we model male lifespan in continuous time using the negative exponential distribution, with a constant baseline mortality rate μ, that corresponds to the genotype with the lowest risk of predation (the yy genotype). Male reproductive success also occurs at a constant baseline rate *R* across a single breeding season, except for the early period of a male’s adult life that occurs in the absence of females. We denote this duration by *T*, the mean degree of protandry in the population (with *T* = 0 indicating no sexual dimorphism in eclosion time). The baseline *R* refers to the rate at which eggs are fertilized by the genotype that most efficiently mates with females (the WW genotype). If we first assume no effects of genotype on mortality or reproduction, lifetime reproductive success is

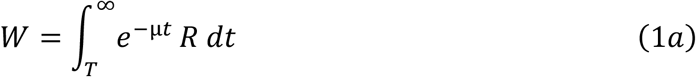

which, after integrating, simplifies to

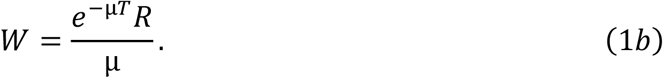

By forming alternative, genotype-specific expressions based on Eq. (1a-b), our framework incorporates simultaneous, interacting effects of a genotype on all relevant fitness components (table 1). This differs from typical models of antagonistic pleiotropy between survival and reproduction, where survival to adulthood is modelled first and only survivors reproduce, and reproductive success is therefore limited solely to those that survived an initial period of predation (e.g. Arnold and Wade 1984; Curtsinger et al. 1994; Rose 1982; Hedrick 1999; Zajitschek and Connallon 2017). Our approach allows for ‘gentler’ impacts where death only removes reproductive events that would have happened at a more advanced age.

**Table 1.** Effects of genotype on male-limited fitness-components.1.

| Trait | Genotype |  |  |
| --- | --- | --- | --- |
|  | yy | Wy | WW |
| Reproductive rate | $(1 - \beta)R$ | $(1 - h_R\beta)R$ | $R$ |
| Mortality rate | $\mu$ | $(1 + h_\mu\gamma)\mu$ | $(1 + \gamma)\mu$ |
| Probability of being alive when females begin emerging | $e^{-\mu T}$ | $e^{-(1+h_\mu\gamma)\mu T}$ | $e^{-(1+\gamma)\mu T}$ |
| Lifespan from female emergence onwards (conditional on being alive at female emergence) | $\frac{1}{\mu}$ | $\frac{1}{(1 + h_\mu\gamma)\mu}$ | $\frac{1}{(1 + \gamma)\mu}$ |
| Expected lifespan spent overlapping with female availability | $\frac{e^{-\mu T}}{\mu}$ | $\frac{e^{-(1+h_\mu\gamma)\mu T}}{(1 + h_\mu\gamma)\mu}$ | $\frac{e^{-(1+\gamma)\mu T}}{(1 + \gamma)\mu}$ |
| Expected lifetime fitness | $\frac{e^{-\mu T}(1 - \beta)R}{\mu}$ | $\frac{e^{-(1+h_\mu\gamma)\mu T}(1 - h_R\beta)R}{(1 + h_\mu\gamma)\mu}$ | $\frac{e^{-(1+\gamma)\mu T}R}{(1 + \gamma)\mu}$ |

To model antagonistic pleiotropy, we modify *μ* and *R* such that expression of the W allele increases male mortality rate, while expression of the y allele decreases the number of eggs fertilized by the male (see table 1). Let *β* and *γ* represent modifications caused by complete expression of the y and W alleles to the baseline reproductive and mortality rates, respectively. *h*_*R*_ and *h*_*μ*_ are the respective dominance coefficients that determine expression in male heterozygotes. When *h*_*R*_ = *h*_*μ*_ = 1, heterozygote males have both a high risk of predation and a low reproductive rate; a ‘worst of both worlds’ deleterious reversal of dominance for polymorphism maintenance. Setting these values to 0.5 indicates additive expression for both traits, while setting them to zero represents complete masking of the negative effects on both traits: this last option represents a ‘best of both worlds’ beneficial reversal of dominance. The genotype-specific fitnesses are

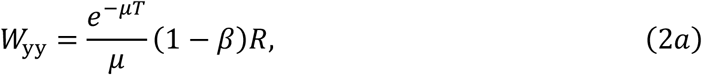

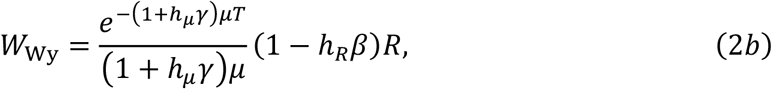

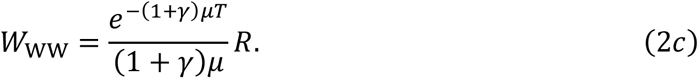

Writing the equations like this neatly partitions the two components of fitness affected by the focal locus. The terms within the fraction relate to each genotype’s effect on mortality. Here, it is useful to remember that the mean of a process with exponentially distributed waiting times is 1 / *λ*, where λ is the rate of occurrence. The fraction therefore represents the mean lifespan of a male (mean waiting time until death occurs), determined by the rate of mortality, but modified by the effect of protandry (if there is no protandry, the numerator reduces to 1 for all genotypes). That protandry modification reduces this quantity to instead reflect the mean length of a male’s *reproductive* lifespan. Everything outside the fraction determines the rate of reproduction during this period, which, in contrast, is not affected by protandry. Following convention (Curtsinger et al. 1994, Hedrick 1999), we use Equations 2a-c to identify conditions where selection maintains balanced polymorphism; that is, where antagonistic pleiotropy produces overdominance for fitness (*W*_yy_ < *W*_Wy_ > *W*_WW_). The condition for balanced polymorphism is

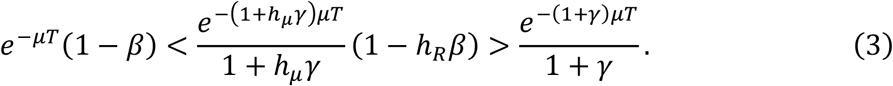

To determine the effects of dominance reversal and protandry, we first explore the potential for balancing selection in the absence of protandry, then show how positive values of protandry change the outcome.

In the absence of protandry, Equation 3 reduces to

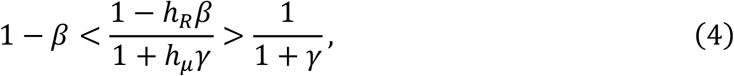

which can be expressed in terms of *β*:

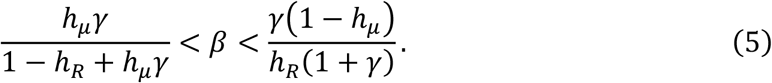

Eq. 5 shows that in the absence of protandry, beneficial reversal of dominance is necessary (though not sufficient) to maintain diversity when there are trade-offs between fitness components that temporally overlap (compare figure 1A with 1B-D). Increasing the strength of dominance reversal, such that the heterozygous genotype increasingly expresses a ‘best of both worlds’ phenotype, broadens the conditions under which polymorphism is predicted. This disproportionately comes at the expense of the parameter space where the W allele is predicted to fix. This effect is so pronounced at high levels of beneficial dominance reversal (e.g. *h*_*R*_, *h*_*μ*_ < 0.2; figure 1C-D) that, in the absence of additional selective forces, conditions where the W allele is predicted to fix are negligible.

**Figure 1.**
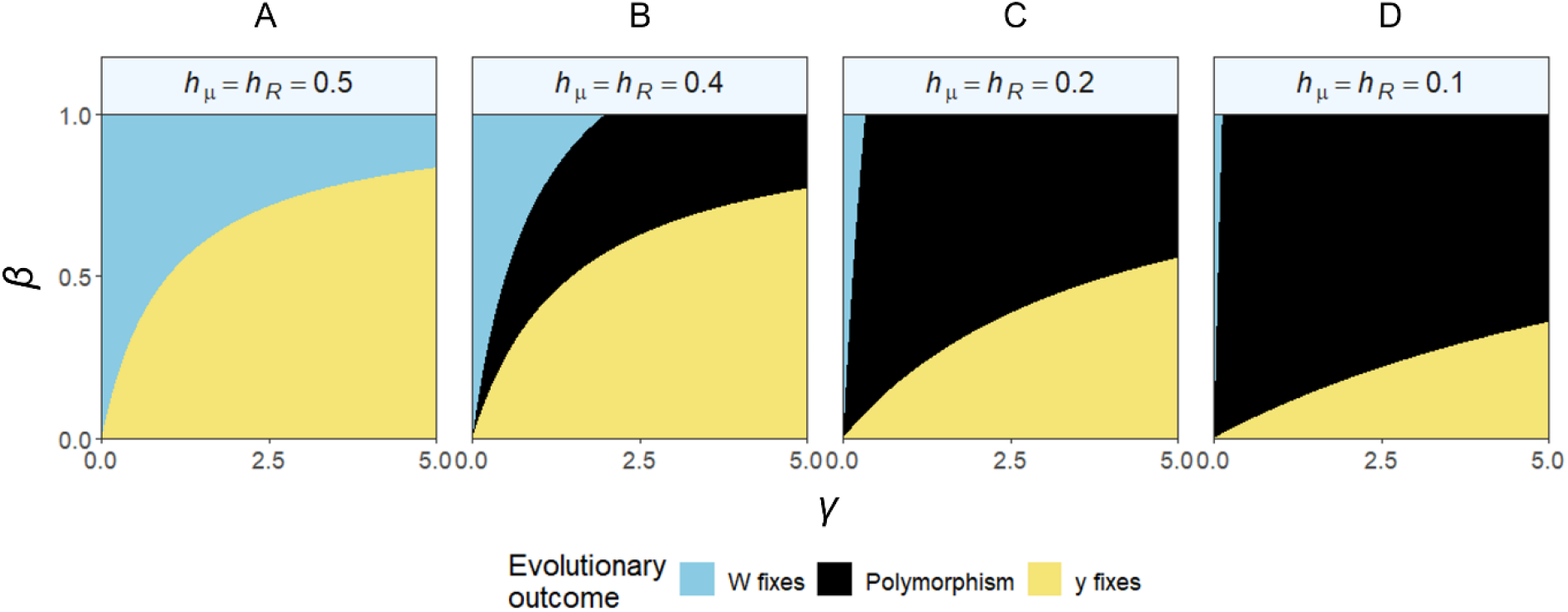
In the absence of protandry, dominance reversals are required to maintain balanced polymorphism. Panels show conditions where the W allele fixes, the y allele fixes, or polymorphism is maintained. *γ* increases the mortality hazard for males carrying the W allele; a value of one doubles the mortality hazard, a value of two triples it, and so on. *β* scales the proportional reduction of the fertilization success of males carrying the y allele, with *β* = 1 removing all reproductive success of males carrying two y alleles (realistically *β* < 1). Dominance coefficients *h*_*R*_ and *h*_*μ*_ control how *γ* and β affect male heterozygotes. Here dominance is equal (*h*_*R*_ = *h*_*μ*_), with the value decreasing across the columns.

Next, we consider protandry. When a male has to survive for some time without appreciable reproductive success (*T* > 0), the reproductive lifespan of a male is no longer equivalent to adult lifespan. An increase in protandry places greater weight on protection from predation for fitness, as a greater proportion of life must be survived before reproduction commences. Intuition suggests that this benefits the y allele, due to the protection it offers against predation, but whether this also impacts the prospects for polymorphism is less clear.

Here, it is informative to evaluate the ratios of *W*_yy_, *W*_Wy_ and *W*_WW_ under biologically relevant values for all parameters: 0 < *h*_*μ*_ < 1, 0 < *h*_*R*_ < 1, 0 < β < 1, γ > 0, μ > 0, and *T* ≥ 0. First, compare the homozygotes’ fitness by evaluating their ratio, *W*_Wy_/*W*_WW_. From eq. (2), this ratio simplifies to

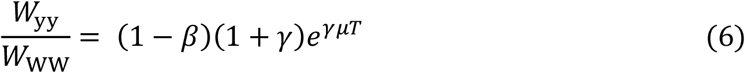

When the right-hand side (RHS) of eq. 6 exceeds 1, W cannot fix, leaving the options open for whether y fixes or there is a polymorphism. A value below 1 predicts the opposite situation where yy cannot fix, thus either W fixes or there is a polymorphism.

To investigate how protandry changes the chances of observing polymorphism, we need to compare to a case without protandry. When *T* = 0, eq. (6) simplifies to

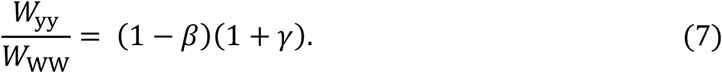

Since biological relevance requires 0 < *β* < 1 (yy’s reproductive rate falls below WW but is not zero), we find that *T* = 0 predicts (1 − *β*)(1 + *γ*) > 0. For different combinations of *β* and *γ*, both (1 − *β*)(1 + *γ*) < 1 and (1 − *β*)(1 + *γ*) > 1 are possible outcomes and thus either homozygote can have higher fitness.

It is relevant to know which homozygote has the higher fitness (eq. 7), as we predict polymorphism only when *W*_Wy_ > *max*{*W*_yy_, *W*_WW_}. Since heterozygote fitness needs to be exceed that of the fitter homozygote, we term the latter its ‘primary competitor’. This proves useful below, though we note that genotypes do not compete the exact same way as alleles do, as e.g. a heterozygote can produce homozygous offspring.

Keeping the above criterion in mind, we now compare the two homozygotes when there is protandry, i.e. *T* > 0.

We first consider the scenario where (1 − *β*)(1 + *γ*) > 1, which can occur if *β* is sufficiently low (to be precise, if *β* < *γ*/(1 + *γ*)). Under these conditions, *W*_yy_ > *W*_WW_ always holds, whether we assume *T* = 0 or *T* > 0. Inspecting eq. 6 reveals that increasing *T* makes the RHS increase from its value at *T* = 0 which already, per assumption, exceeded 1. Thus, when (1 − *β*)(1 + *γ*) > 1, the primary competitor of the heterozygote is yy, regardless of the value of *T*. Note that yy suffers little in terms of its reproduction when β is low, making the result intuitive.

Next, we consider what happens in the reverse situation where (1 − *β*)(1 + *γ*) < 1. Now, when *T* is low, the RHS of eq. (6) remains smaller than 1, and *W*_WW_ > *W*_yy_. But if *T* is sufficiently large 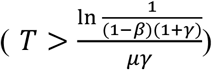, the homozygote fitnesses flip to *W*_yy_ > *W*_WW_. In other words, long protandry starts favoring the morph with the survival advantage, and whether *W*_yy_ or *W*_WW_ is the heterozygote’s primary competitor depends on *T*.

We next show that (a) if the primary competitor is WW, then increasing *T* improves the prospects for polymorphism, and (b) if the primary competitor is yy, then increasing *T* harms the prospects for polymorphism.

Let us proceed with (a) first. From the above, we know that WW is the heterozygote’s primary competitor if the situation combines (1 − *β*)(1 + *γ*) < 1 with 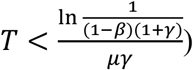. In this case, we are interested in knowing whether *W*_*Wy*_/*W*_*WW*_ increases or decreases with the value ot *T*. This ratio takes the form

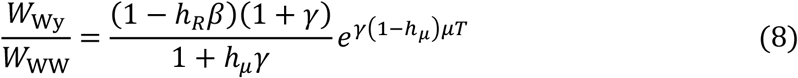

Within our biologically relevant set of parameter values, the ratio 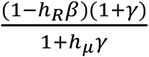 is always positive and does not depend on *T*. Thus, whether the RHS of eq. 8 increases with *T* depends solely on the term 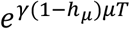. This term is an increasing function of *T* and thus the entire ratio *W*_Wy_/*W*_WW_ can only increase with *T*, or, in the special case of *h*_*μ*_ = 1, remain unchanged. The conclusion is that when Wy and WW are competing, protandry can only ever favor Wy. Protandry can thus shift the system from “W fixes” to a polymorphic state; it can never achieve the opposite. Note though the possibility that the outcome remains unchanged, which happens if the RHS increases with *T* without crossing the value 1, or does not depend on *T* at all (the special case *h*_*μ*_ = 1).

Next, consider statement (b), i.e. assume that the heterozygote’s primary competitor is yy. This happens when the set of conditions at the beginning of the (a) section is not fulfilled. Now, the ratio of interest is

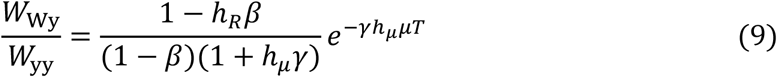

The term 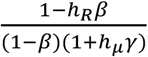 is, for our set of biologically realistic parameter values, always positive, and once again the exponential term determines whether the ratio increases or decreases with *T*. The term 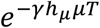 clearly decreases with *T* (again we use information on our biologically relevant parameter values), and thus, whenever protandry is mediating the competition between yy and Wy, increasing protandry can only shift the solution from polymorphism to yy, never the opposite.

A graphical view of the conditions for polymorphism shows that increasing protandry (moving along the *x* axis of each subplot in figure 2) never moves the system from a yellow-fixing prediction to polymorphism, while if the starting point is one where the fixation of the white allele is predicted (high *β*), it is possible to move to a polymorphism when protandry increases. Thus, polymorphism replaces regions that, without protandry, would favor fixation of the W allele. As protandry increases further, polymorphism can also disappear again, replaced by the fixation of y. It is noteworthy that the examples in figure 2 use a dominance set that is equal and additive (*h*_*R*_ = *h*_*μ*_ = 0.5), which are conditions that would, in the absence of protandry (fig. 1A), never produce polymorphism.

**Figure 2.**
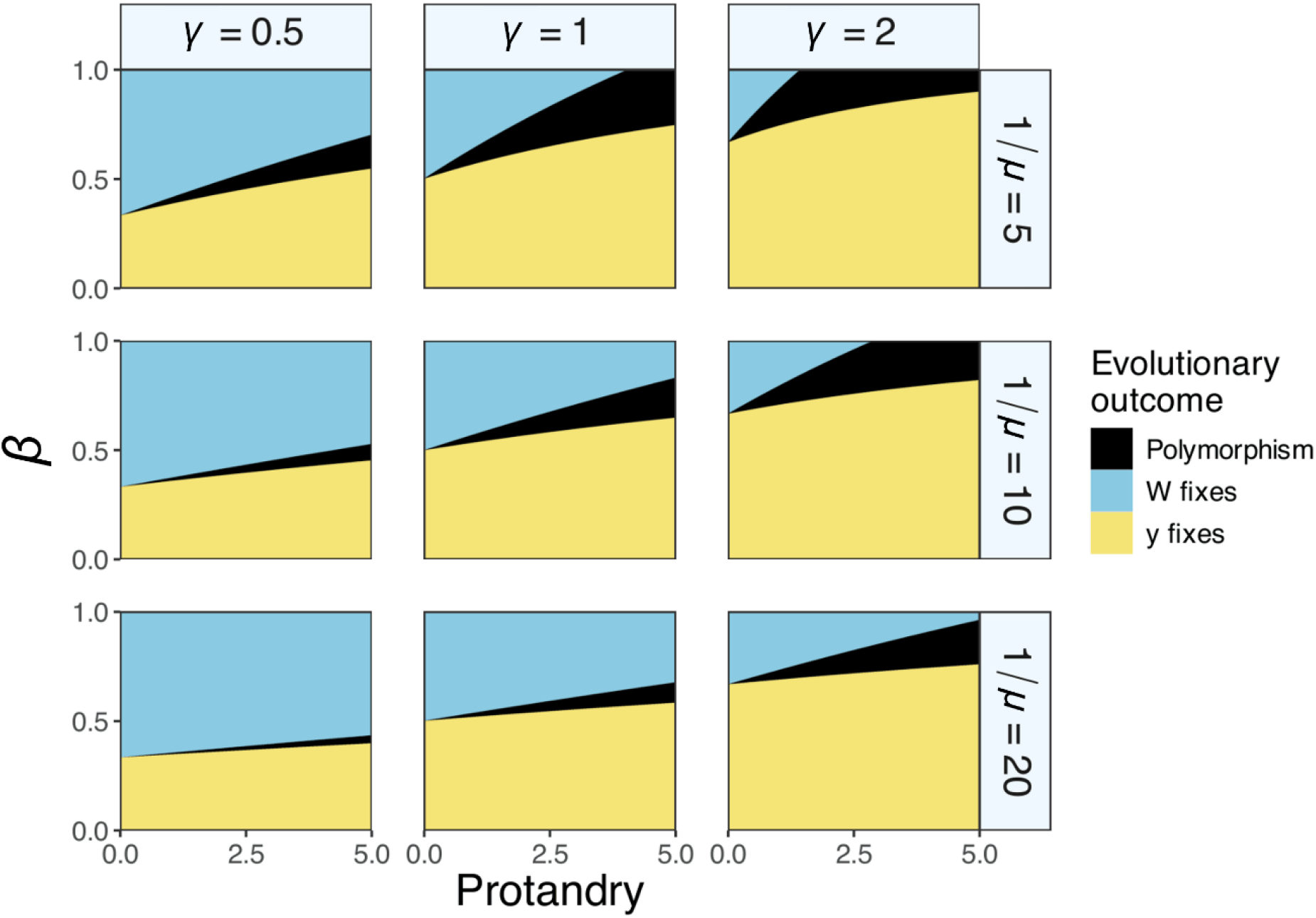
Protandry can produce balancing selection in the absence of beneficial reversal of dominance. Subplots differ in male lifespan (rows) and γ, and all use additive dominance values *h*_*R*_ = *h*_*μ*_ = 0.5. The length of protandry (*T*) is varied on the x-axis; the time unit should be interpreted with the knowledge that the mean lifespan of the longest-lived male genotype is 1/*μ*. The fixation of either W or y at zero protandry is sufficient to determine whether increasing protandry can shift the solution to a polymorphic state or not.

### Individual-based model: Methods

Our analytical model above achieved a very general prediction for the effect of protandry on polymorphism maintenance under antagonistic pleiotropy, but the main prediction — whether protandry helps or harms depends on which homozygote is more fit — was derived under heavily simplified assumptions that are worth relaxing. Specifically, the analytical model assumes two pulses of adult emergence (first males, then females), and that each male accumulates matings at a constant rate commencing at female emergence and continuing until the death of the male. A more realistic model, which we build below, uses mate encounter rates that depend on mate searching capabilities and the densities of receptive males and females, which change dynamically throughout the breeding season due to mortality and post-mating refractory periods. Our individual-based model is parameterized with estimates based on experimental results from our model system.

The simulations track allele frequencies at a diploid color locus in a finite population comprised of two sexes, with non-overlapping generations but modelled in continuous time within a season. As in the analytical model, we consider two alleles y and W that produce diploid genotypes with discrete wing-color morphs in adult males. We assume the alleles are not expressed and are thus selectively neutral in females. The mating system is monandrous and polygynous, such that females mate once and males can mate multiply. After the population is initially seeded with alleles, we assume no mutation or migration and strict Mendelian inheritance, such that allele frequency changes are limited to the effects of drift and selection.

#### Within-generation dynamics

While generations are discrete, we track within-generation dynamics in continuous time, using a Gillespie algorithm (Gillespie 1976). Specifically, individuals move between states depending on four types of events that occur within a breeding season: emergence into the mating pool, mating, post-mating recovery and death.

Each event occurs at a specified rate (see below for details), and the algorithm selects one to happen among several events that are ‘competing to happen’ (see Kokko 2024 for an extended explanation of the method). Time therefore does not progress in predefined discrete steps but takes any real-numbered value according to when each event takes place. We track individuals’ changing states and accumulated matings, until no further matings are possible: either because all females have mated or because there are no females or males alive. While the Gillespie algorithm allows time to be measured in any units, we chose units such that the passage of one unit of time is equivalent to one day in the Finnish breeding season of the wood tiger moth.

Below, we list the event types separately, but note that since a Gillespie algorithm does not use time steps of fixed duration, the algorithm moves flexibly between the event types: the soonest event to occur is executed first and the choice between events depends on the rate-specific waiting time.

#### Emergence

Individuals enter the mating pool upon emergence from their pupae, after which we consider them sexually mature and exposed to predation. Emergence times are handled as ‘time stamps’ (*see* Kokko 2024) that are pre-allocated to all individuals. Emergence is chosen as the next event, when an individual’s time stamp is smaller than the time at which any other event occurs. Emergence times are drawn from the sex-specific probability distributions

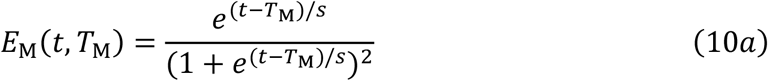

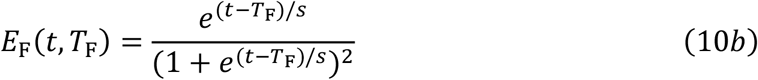

where *E*_M_(*t, T*_M_) and *E*_F_(*t, T*_F_) are the numbers of males and females emerging at time *t*, respectively (as in Zonneveld and Metz 1991). *T*_M_ and *T*_F_ determine the mean emergence time for males and females respectively, and the parameter *s* determines the variance – effectively changing the length of the possible emergence period. We assume the same variance for both sexes and that the color locus has no effect on emergence time in either sex.

We vary the degree of protandry by changing *T*_F_ while keeping *T*_M_ constant. Importantly, this means that protandry does not coevolve with hind-wing coloration in our simulations (see Discussion). We set *T*_M_ = 0 (note that the absolute value of *T*_M_ has no biological meaning) and vary *T*_F_ to be integers between 0 and 5, yielding a corresponding degree of protandry between 0 and 5 days.

#### Mating and death

Only individuals that have emerged can be chosen to mate or to die. We model genotype-specific male mating success by varying the rates, *v*_*ij*_, at which each male genotype (*i* and *j* represent alleles; each can be W or y) locates receptive females. The rates for homozygote males were estimated based on morph-specific mating success reported in Selenius et al. (2025) (table 2, see Supplementary Methods for details). Note, that while we conceptualize these rates as the speed at which males locate females, they can also include observed differences in mating success resulting from mate choice.

**Table 2.** Values for the set of parameters used in our individual-based simulations (see Supplementary Methods). *h*_*R*_ and *h*_*μ*_ are the dominance coefficients for male search and mortality rates, respectively, which we let vary independently from 0 to 1.

| Parameter | Value |
| --- | --- |
| Protandry | $T$ |
| WW mate search rate ($v_{WW}$) | 0.34 |
| yy mate search rate ($v_{yy}$) | 0.18 |
| Wy mate search rate | $h_R v_{yy} + (1 - h_R) v_{WW}$ |
| WW mortality rate ($\mu_{WW}$) | 0.18 |
| yy mortality rate ($\mu_{yy}$) | 0.1 |
| Wy mortality rate | $h_\mu \mu_{WW} + (1 - h_\mu) \mu_{yy}$ |
| Female mortality rate ($\mu_F$) | 0.09 |
| Male refractory period ($r$) | 1 |

Following the mass-action law (Hutchinson and Waser 2007; Kokko 2024), let *F*_*t*_ be the number of adult virgin females alive at time *t, M*_*ij,t*_ the number of males in the mating pool with genotype *ij* at time *t*, and *v*_*ij*_ the rate at which males of genotype *ij* search for females. Across the population, the total rate at which an event of the type ‘a genotype *ij* male mates’ occurs is *v*_*ij*_*M*_*ij,t*_*F*_*t*_. The wait time to the next event of this type is thus exponentially distributed with parameter *v*_*ij*_*M*_*ij,t*_*F*_*t*_; if sampling from this distribution yields a time that is sooner than any other type of event, we choose one male from the appropriate category and one female to form the mated pair (female genotypes were sampled from the current frequency of each genotype irrespective of the time within a season, as female survival does not depend on genotype). Mating causes the female to permanently leave the mating pool, while a male temporarily exits the mating pool for a time of length *r* (‘refractory period’, table 2), which includes the time required for mating to complete and for males to begin searching again.

We parameterised the male refractory period by loosely estimating it from mating duration data in *A. plantaginis* (Santostefano et al. 2018)(table 2, see Supplementary methods for details). The refractory period was kept the same for each mating, thus return to the mating pool uses time stamps (*sensu* Kokko 2024), not exponential distributions.

After an individual has emerged, it faces a constant risk of mortality. For females, mortality occurs at a genotype-independent rate *μ*_F_. In contrast, to capture the effect of wing coloration on predation risk, we let the male mortality rate vary with genotype, such that *μ*_*ij*_ represents the mortality rate for a male with genotype *ij*. We assumed that predators were equally likely to target males that were in or out of the mating pool, such that the mortality rate was unaffected by an individual’s current state (conditional upon it being alive). At time *t*, the total rate at which a death of a female occurs is *μ*_*F*_*F*_*t*_ (we track this only for females still available as mates, as post-mating mortality does not affect the evolutionary outcome), while for males of a specific genotype this equals *μ*_*ij*_*M*_*ij,t*_. The mortality rates we used (table 2) were informed by a combination of laboratory longevity studies and field predation experiments for both males (Nokelainen et al. 2012) and females (Lindstedt et al. 2011) (see Supplementary Methods).

#### Between generations

Upon completion of a breeding season, mated females produce offspring in the Mendelian ratios that correspond to their genotype and that of their mate. We assume that selection is soft and that density regulation occurs implicitly and unbiasedly with respect to genotype during the juvenile life stage, such that while juvenile population sizes greatly exceed that of adults, the number of individuals surviving to adult emergence is set to 2000 individuals in each generation. This means we do not explicitly model juveniles, instead we sample 2000 new recruits to emerge and form the next generation. We first generate 1000 male recruits (following an unbiased sex ratio). Each recruit’s mother is randomly chosen from among the mated females of the parental generation, and the mother’s genotype together with the genotype of the associated male is used to yield the recruit’s genotype based on Mendelian inheritance. The remaining 1000 recruits are assigned as females, with a genotype distribution identical to that of males.

We ran the simulation for different combinations of protandry *T*, male heterozygote mate searching rates (*v*_Wy_) and mortality rates (*μ*_Wy_). The latter two varied as a function of the dominance parameters (table 2), where *h*_*R*_ < 0.5 and *h*_*μ*_ < 0.5 imply that heterozygotes resemble the better performing homozygote more than the alternative homozygote (beneficial reversal of dominance). Every combination was run for 1000 generations three times: once with the W allele starting at frequency *p* = 0.5 (unbiased), once with *p* = 0.8 (W-biased) and once with *p* = 0.2 (y-biased). The first generation was assumed to be in Hardy-Weinberg equilibrium.

Additionally, to observe if heterozygote advantage is indeed the primary mechanism behind protected polymorphism in our model, we estimated the relative fitness of each genotype by calculating their relative reproductive success and taking its geometric mean across ten generations (Haldane and Jayakar 1963). As estimating becomes impossible late in a simulation run in cases where one morph has fixed, we chose to estimate fitness early, using data from generations 11 to 20.

### Individual-based model: Results

Using empirical estimates for mating and mortality rates, we find that the strength of beneficial dominance reversal required for stable polymorphism depends on the degree of protandry. Protandry also affects which of the alleles fixes when polymorphism isn’t maintained.

Our results from the individual-based model (fig. 3) replicated the analytical model’s main finding. Recall that the prediction is that if the heterozygote’s primary competitor is WW, then increasing protandry can switch the system from WW to a polymorphism; when, however, yy is the primary competitor, then protandry cannot promote polymorphism. This is clearly visible in the sequence of results (fig. 3) where we investigate the effect of dominance (x and y axes), while increasing the amount of protandry (subplots). In figure 3A-3D, the primary competitor is WW, as this morph fixes as soon as the dominance combination no longer permits the maintenance of a polymorphism. Here, increasing protandry clearly helps the maintenance of polymorphism, visible as the dominance parameter region permissive of a polymorphism increasing from figure 3A to figure 3E. Phrased another way, when protandry increases, the strength of beneficial dominance reversal required to maintain polymorphism decreases until polymorphism is also maintained at additive dominance (fig. 3E, where *h*_*R*_ = *h*_*μ*_ = 0.5). Increasing protandry further (fig. 3E-3F) enhances the importance of survival so that yellow now beats white and is thus the heterozygote’s primary competitor. Accordingly, protandry now has an opposite effect on polymorphism: the prediction that ever-longer protandry should shrink the parameter range for polymorphism (as shown by the analytical model) is also shown in the individual-based model (the region of polymorphism shrinks from 3E-3F).

**Figure 3.**
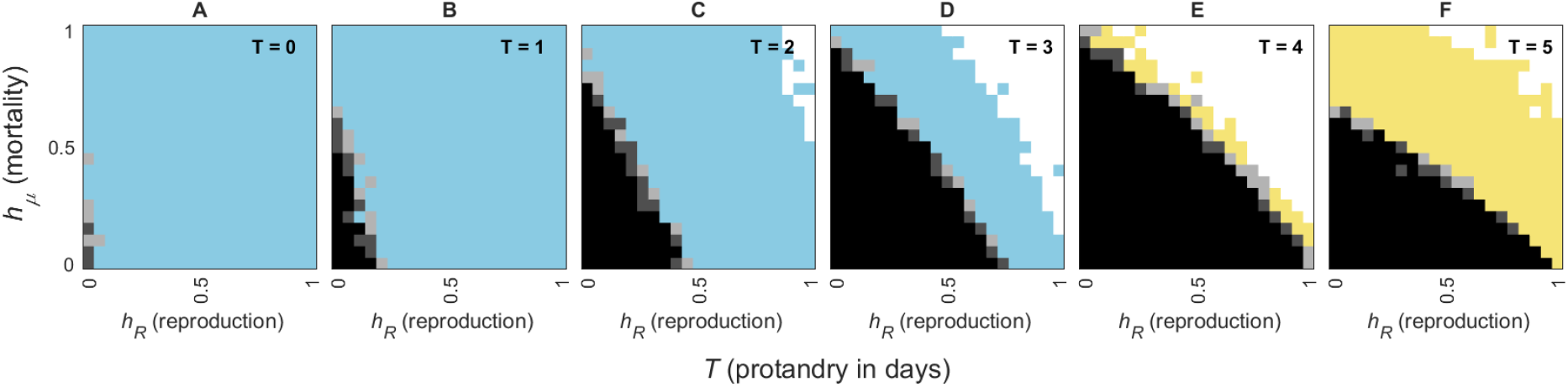
Polymorphism across dominance parameters and different degrees of protandry measured in days between mean male and female emergence after 1000 generations of selection. The black squares represent regions of polymorphism across all three starting frequencies, and the shades of grey represent those regions of polymorphisms that occurred only in two (dark grey) or one (light grey) of the three starting frequencies. Blue and yellow squares represent populations where the W or y allele respectively fixed across all starting frequencies, whereas white squares represent populations where a different allele fixed under different starting frequencies due to stochastic effects. The bottom left corner, where most polymorphism is found, shows beneficial reversal of dominance. Parameter values that were not varied were taken from table 2.

The results show that requirements for the strength of dominance reversal are mildest, and the parameter region for polymorphism the most permissive, when the two homozygotes are roughly equally fit, which happens at intermediate protandry. As a whole, the individual-based simulation indicates that the results from the analytical model extend to a much more realistic setting of finite mate finding rates, males interfering with each other’s mating success by making females unavailable as soon as they have mated, and imperfect temporal separation of the male and female emergence distributions.

Protandry also changes the details of the shape of the dominance parameter space where polymorphism is maintained (fig. 3). At low protandry, polymorphism is maintained only when heterozygotes have very high search rates (i.e., similar to the homozygote with high mating success), while at high protandry maintenance of polymorphism occurs only when heterozygote mortality is appropriately low (i.e. very similar to the homozygote with low mortality). This matches the intuition of mortality being emphasized more in the fitness equation when females emerge significantly later than males.

The regions of polymorphism across the dominance parameters match well with observed heterozygote advantage (fig. S2), further supporting it as the primary mechanism maintaining polymorphism in the species under antagonistic pleiotropy.

## Discussion

Previous theoretical works on antagonistic pleiotropy have provided us with general conditions under which protected polymorphism can occur (Curtsinger et al. 1994; Hedrick 1999). The conditions of strong selection, relatively equal selection coefficients, and dominance reversal provide a solid footing for polymorphism, but these statements are not informative regarding how such conditions emerge in nature, or whether they are biologically realistic. The classic way of modelling fitness components sequentially (e.g. survival first and then reproduction of the survivors) does not take into account features of realistic life cycles, where lifespans may be cut short by predation after some offspring production has already taken place; this necessitates a more detailed temporal look at the dynamics within a lifespan.

Here, we demonstrate that relevant differences in fitness can arise from morph-specific variation in mate searching abilities along with temporal differences in male and female emergence, combined with differences in vulnerability to predation. By applying a population genetic model, we show how the temporal dynamics of fitness accumulation can result in a protected polymorphism. While the analytical model makes some stark simplifications of male reproduction, our individual-based simulations parameterized by data from a natural system show the same result: modest protandry (a few days) can help maintain a polymorphism while males needing to survive for longer durations before female availability peaks make the best-surviving male morph fix. This provides a solid theoretical footing for the idea that antagonistic pleiotropy underlies the maintenance of a sex-limited polymorphism in *A. plantaginis*.

Our population genetic model shows that protandry can increase the likelihood of polymorphism only when the primary competitor of the heterozygotes in the absence of protandry is the morph with the mating advantage (WW). While adult moths are under a continuous predation pressure by mostly avian predators (see e.g. Rönkä et al. 2020; Nokelainen et al. 2012; 2014), we expect strong sexual selection on males’ ability to locate females in the moth mating system, where females are sparsely distributed and the reproductive period is short (Herberstein et al. 2017). Sexual selection therefore determines the evolutionary outcome when females are readily available shortly after male emergence, but as protandry increases beyond a few days – as determined by our simulations – the morph-specific differences in survival start to weigh more and drive the fixation of the y allele.

The four days difference in male and female emergence, which provides the optimal scenario for polymorphism, is within realistic estimates for protandry in the wood tiger moth. Overall, protandry is common across *Lepidoptera*, where it is likely under selection in mating systems where males benefit from earlier access to females (Zonneveld 1992; Singer 1982; Fagerström and Wiklund 1982; Ekrem and Kokko 2023). Additionally, protandry can emerge as a byproduct of sexual size dimorphism and correlated differences in developmental time (Teder 2014; Teder et al. 2021). This may be the case in the wood tiger moth, where females are larger than males (see data from De Pasqual et al. 2022). Direct evidence for protandry comes from rearing experiments by Ojala et al. (2005), which suggest the difference in male and female developmental time in the wood tiger moth is approximately three days across different larval diets – although there is large variation between diets. In addition, capture-mark-recapture data from a Finnish population of wood tiger moths confirms earlier emergence of males and females in nature (Gordon et al. 2026).

Because we did not aim to produce a model for the evolution of protandry *per se* (many models exist already, especially for Lepidopterans, e.g. Ekrem and Kokko 2023; Degen et al. 2015; Iwasa et al. 1983; Iwasa and Haccou 1994; Fagerström and Wiklund 1982; Wiklund and Fagerström 1977; Kubo et al. 2025; Zonneveld and Metz 1991), we assumed that protandry is selected for similarly across all males. One could conceivably argue that if males are able to adjust their emergence in a genotype-specific way, WW and Wy males should evolve to emerge later than yy males, as the former type lacks the antipredatory coloration that permits survival through a prolonged period of time before females arrive. This outcome would make the relevant value of *T* morph-specific in the fitness equations. The reason that we did not pursue this route is that pedigree data suggests no difference in male emerge timing when males are grouped phenotypically (white and yellow, table S1). It is possible that the genetic conflict over timing is simply unresolved in males, which is perhaps to be expected for a trait controlled by a single locus (Nokelainen et al. 2022; Brien et al. 2023) and not by a supergene (Chouteau et al. 2017; Thomas et al. 2008). Pleiotropic effects (Andrade et al. 2019) could, of course, occur, but we have no evidence for them in our case.

Our results align well with existing general work on balancing selection: we showed it can maintain color polymorphism under additive dominance under highly specific conditions, while any form of beneficial reversal of dominance greatly increases the likelihood of polymorphism (as also shown by Hedrick 1999; Kidwell et al. 1977). Therefore, realistically, we should expect some degree of dominance reversal in the wood tiger moth for the polymorphism to be maintained. Beneficial reversal of dominance has been traditionally considered to be exceedingly rare (R. F. Hoekstra et al. 1985; Curtsinger et al. 1994), but recent empirical findings suggest that it is not only plausible but can be found across several clades (Gautier et al. 2018; Jardine et al. 2021; Johnston et al. 2013; Karageorgi et al. 2025), especially in the case of sex-specific dominance (Barson et al. 2015; Connallon and Chenoweth 2019; Grieshop and Arnqvist 2018). While the evolution and genetic mechanisms of dominance reversal still remain under discussion (Grieshop et al. 2024; Flintham 2025), heterozygote fitness may also resemble that of the fitter homozygote across contexts without the need for specific expression modifiers due to ecology (see e.g. Johnston et al. 2013), which presents a more likely scenario for the fitness trade-off between color morphs in the wood tiger moth.

As dominance reversal requires the heterozygotes to resemble the best morph both in terms of mating success (WW) and survival (yy), we must consider potential mechanisms underlying morph-specific fitness advantages. The precise causality behind the WW males’ higher mating success is unknown, but *yellow* family genes have been linked to male courtship behavior with varying effects between species (Connahs et al. 2022; Massey et al. 2019). Alternatively, differences in mating success may result from greater flight activity of the white males compared to yy males (Rojas et al. 2015), or other costs associated with production of the pheomelanin-based yellow pigment (Galván and Møller 2011) – costs that phenotypically white Wy males can likely avoid. Empirical data in any case indicates that the Wy morph’s mating success is similar to that of the WW morph (Selenius et al. 2025), which is beneficial for the polymorphism.

The survival advantage of the yy males is more straightforward to understand. The yellow hindwing coloration is a classic aposematic warning signal, shown to both aid predator learning and cause an innate avoidance reaction in avian predators (Gittleman and Harvey 1980; Lindström et al. 1999; Schuler and Hesse 1985). While heterozygotes resemble the WW morph to the human eye, relevant predators can likely distinguish between WW and Wy due to differences in ultraviolet reflectance (Nokelainen et al. 2012). The empirical support for the effective usage of UV coloration as part of an aposematic signal is inconclusive (Lyytinen et al. 2001, 1999; Crowell et al. 2024; Corral-Lopez et al. 2021), and all previous publications on predation on the wood tiger moth color morphs have grouped WW and Wy males together (see e.g. Nokelainen et al. 2012; 2014; Rönkä et al. 2020). Since empirical evidence suggests that increased chromatic contrast benefits males even within the white phenotype (WW and Wy) (Nokelainen et al. 2012), testing whether strong UV reflectance is an antipredator trait prolonging the expected lifespan of Wy males is a clear avenue for further empirical work.

Moving beyond selection coefficients that are applied ‘once per generation’, and deriving fitness explicitly based on events that can occur during an individual’s life, can yield much insight into the mechanisms that generate relevant differences in fitness. Interestingly, a recent study on a female-limited color polymorphism in *Colias* butterflies (Klein et al. 2025) shows that timing differences between males and female morphs (in this species males emerge first, followed by white females and, last, orange females) is potentially key to understanding the polymorphism in that system too; though in their case the causative agent appears to be competition for resources provided by males, and not related to predation. Taken together, the diversity of potential mechanisms suggests that timing traits – protandry among them – may be important facilitators of color polymorphism across *Lepidoptera*. More broadly, our results reinforce the value of embedding species-specific ecology into models of antagonistic pleiotropy. Similar approaches have yielded insights in other systems (e.g. *Microbotryum violaceum*: Tellier et al. 2007; annual plants: Brown and Kelly 2018), highlighting the wide range of ecological scenarios capable of generating protected polymorphisms.

## Notes

### Competing Interest Statement

The authors have declared no competing interest.

### Funding

- Research Council of Finland: 135595
- Alexander von Humboldt Foundation
- Deutsche Forschungsgemeinschaft: 407023052

