## Supplementary Materials for "Beyond abstract selection coefficients: Protandry impacts the buildup of heterozygote advantage over the lifespan in a color polymorphic moth"

* Shared first authorship

^1^ Organismal and Evolutionary Biology Research Programme, Faculty of Biological and Environmental Sciences, University of Helsinki, Viikinkaari 1, 00790, Helsinki Finland.

^2^ Institute of Organismic and Molecular Evolution (iomE), Johannes Gutenberg University of Mainz, Mainz, Germany.

^3^ BioFrontiers Institute, University of Colorado Boulder, Boulder, Colorado, USA.

^4^ Institute for Quantitative and Computational Biosciences (IQCB), Johannes Gutenberg University of Mainz, Mainz, Germany.

**Supplementary Methods**

**Parameterizing the individual based simulations**

The simulation presented in the main text is designed to match the life history of *Arctia plantiginis*. To do this, we used field data collected from the northern range of the species, combined with controlled, laboratory experiments. These studies tend to produce estimates in the form of probabilities (bounded between 0 and 1), e.g., for the likelihood of a mating occurring over a predefined timespan. Our individual-based simulation requires estimation of various rates (above 0 but not bound above), e.g., a high mating rate means that the probability that a mating has *not* happened yet declines rapidly over time. This requires transforming probabilities to rates, which was done using the following procedure.

Let *p*(*t*) denote the cumulative probability that an event (e.g., death, or a mating) has occurred by time *t*. Its complement

$$\begin{aligned} q\left( t \right)= 1-p(t)=e^{-\lambda t}\#\left( S1 \right) \end{aligned}$$

indicates the probability that the event has not yet happened when the individual has been observed for *t* time units. Solving equation *S*1 for λ yields the rate at which this event occurs:

$$\begin{aligned} \lambda=-\frac{\text{ln}\left( 1-p(t) \right)}{t}\#\left( S2 \right) \end{aligned}$$

Note the biologically plausible minus sign: the logarithmic function yields negative values when 0 < *p*(*t*) < 1, and λ as a whole remains positive (assuming observation time *t* > 0).

*Male mating rates*

We used data from small cage mating experiments presented in Selenius et al. (2025) to approximate the rate at which *A. plantaginis* males of each genotype mate with females. These data were generated by competitive mating experiments, where virgin females were presented with males of two of the three yellow-e genotypes, replicated for every possible female-male-male genotype combination. The variable of interest here is the probability that a male mates with the cohabiting female within a single evening. We make the simplifying assumption that mating success measured across one night is an accurate proxy of mating success across the breeding season.

To estimate the probability of nightly mating success, we used a generalised linear mixed model (GLMM), with mating success as response variable and male colour genotype as the predictor variable. We also included generation and year as random effects to control for variation in mating success across the experimental period (see Selenius et al., 2025 for details on variables). Since the GLMM uses a logistic link function to transform the binomial response variable, we transformed the model estimates back into mating probabilities (*p*) using the following equation

$$\begin{aligned} p=\frac{e^{\alpha+\beta}}{1+e^{\alpha+\beta}}\#(S3) \end{aligned}$$

where *α* is the model intercept and *β* is the regression coefficient for our genotype of interest. By setting the WW genotype as the intercept, we can solve eq. S3 for both WW and yy genotypes when the regression coefficient is either zero or the estimate for yy, respectively. These probabilities represent the likelihood of mating after *t =* 1 (as unit of *t* is one day in the Finnish breeding season), and substituting these values into eq. S2 yields the rates used in our simulations.

*Mortality*

We estimated mortality rates for both male homozygotes (WW and yy) by combining empirical data from two previous publications. To find the mortality hazard due to intrinsic causes, we used data on longevity of laboratory stock individuals collected during the mating experiments conducted in Selenius et al. (2025). These data report lifespan for both sexes, measured under standard rearing conditions (see De Pasqual et al., 2022; Nokelainen et al., 2022 for a more detailed description of rearing protocol). Most importantly for our purposes, these conditions lack all forms of predation and thus provide an estimate of intrinsic mortality. To estimate the rate of mortality from lifespan data, we calculated median male lifespan: the number of days elapsed by which 50% of the cohort (here, all males) perished. Put another way, this is the time *t* where *p* = 0.5, where *p* denotes the probability of death. While these data only separate males by human-observable phenotype (white and yellow), we found no difference between the two and therefore used the median of the entire male cohort data to estimate *t* when *p* = 0.5. Substituting these values into Eq. S2 gives the baseline mortality rate for all male genotypes.

To generate estimates of morph-specific predation rates, we used the data from Nokelainen et al. (2012), who recorded the proportion of attacked pinned moths in the wild for the white and yellow phenotype. The proportion of pinned moths removed by predators after 5 days was 0.436 and 0.221 for the white and yellow morphs, respectively. We used the proportion of predated yellow moths as the probability of death for the yy morph after *t* = 5 days and substituted these values into eq. S2 to yield a rate estimate for yy males. As the homozygote (WW) and heterozygote (Wy) individuals were pooled as the ‘white morph’ in Nokelainen et al. (2012), we lack a direct way to estimate the WW mortality rate separately from Wy. We therefore base our values on the graphically reported differences in hindwing chromatic contrast between the two white male morphs and their effect on probability of survival (see fig. 2b in Nokelainen et al., 2012). Since Wy males have generally higher chromatic contrast than WW males (Nokelainen et al., 2022), we compare the associated predicted probabilities of survival, and arrive at an estimated difference in the probability of death between the white genotypes ($p_{\mathrm{WW}}/p_{\mathrm{Wy}}$) of 1.125. Using this value, and assuming an approximate 1:1 ratio of the white genotypes within the experiment, we arrived at a corrected probability of death after *t =* 5 days for the WW homozygotes of 0.462. We substituted these values into eq. S2 for a rate estimate for WW males. Total genotype-specific mortality rates were calculated as the sum of the intrinsic (senescence) and extrinsic (predation) mortalities (note that additivity is correct when interpreting mortalities as rates).

Female mortality rate was also estimated with the same method using a combination of longevity data (same dataset as for males, Selenius et al., 2025) and predation experiment results from Lindstedt et al. (2011), who recorded the attacks by avian predators on pinned female moths in nature. Similar to the predation experiment on males (Nokelainen et al., 2012), this experiment also tracked the proportion of females removed by predators after 5 days (Lindstedt et al., 2011). While a previous publication has suggested that females have significantly shorter lifespans than males (Gordon et al. 2015), we did not find evidence for this in these datasets.

*Male refractory period*

After mating, *A. plantiginis* males guard females for an extended duration, temporarily removing them from the mating pool. We estimated the length of this mate guarding refractory period using mating duration data presented in Santostefano et al. (2018). We estimate this parameter as a duration rather than a rate, since a rate interpretation is not biologically plausible for processes that require completing a finite task (see Kokko 2024: predation can be modelled as a rate since predator striking capabilities are typically equivalent regardless of how long a prey item has been available, while the capacity to mate is often diminished immediately after a mating event but increases with time post-mating). We estimated the duration by taking the median value of all mating durations, irrespective of male genotype. We note that this estimate should be treated with caution, as high variance and the lack of estimates on other relevant traits such as sperm production make it difficult to estimate a precise refractory period for males.

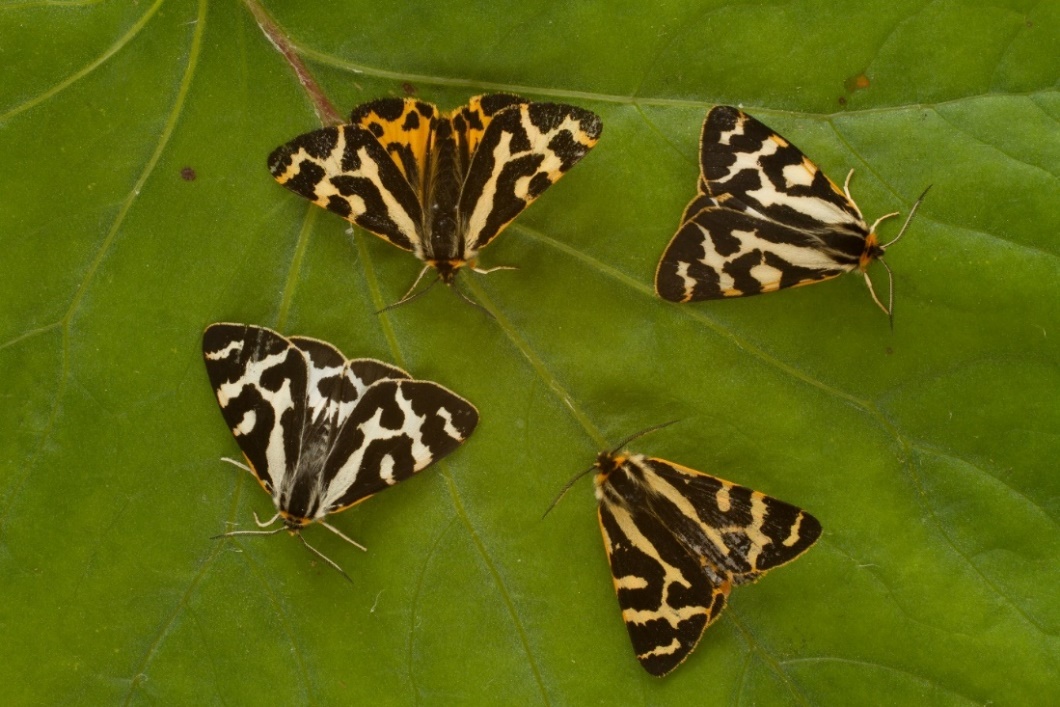

**Figure S1**. Four wood tiger moth males representing the three color morphs: WW (bottom left), Wy (top right) and yy (top left and bottom right). Both WW and Wy males appear phenotypically similar with white hindwings.

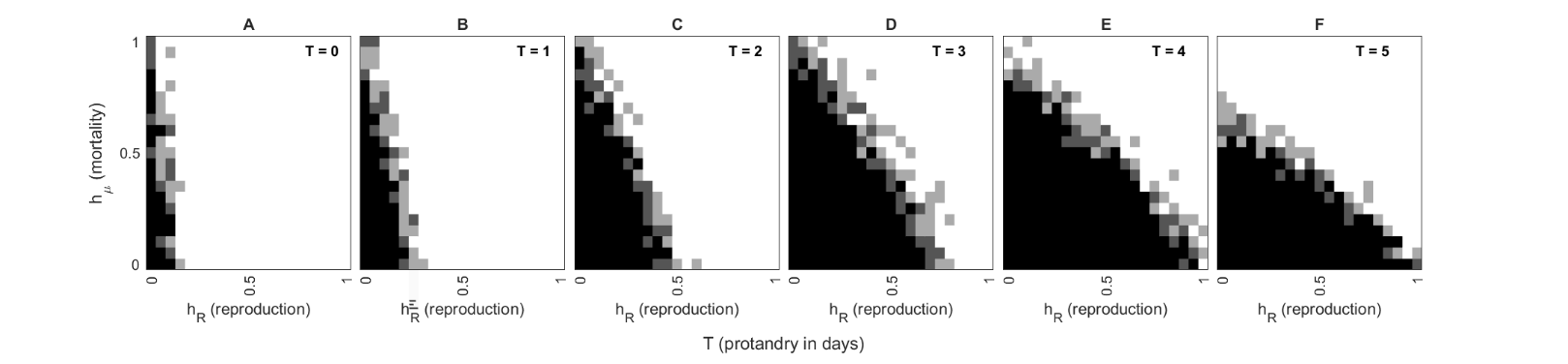

**Figure S2**. Heterozygote advantage across dominance parameters (x and y axis) and degrees of protandry (subplots a-f). The heterozygote advantage was calculated through the geometric mean of relative fitness across 10 generations (generations 11-20). We find regions of heterozygote advantage to match regions of polymorphism across the dominance parameter space. Black squares indicate heterozygote advantage across all three starting frequencies, and dark and light grey indicate heterozygote advantage over two or one of the starting frequencies. Across all panels, *v*_WW_ = 0.34, *v*_yy_ = 0.18, *μ*_WW_ = 0.18, *μ*_yy_ = 0.1 and *N* = 1000.

**Table S1**. Average, standard error and standard deviation of male developmental time, separated by the color phenotype (white including both WW and Wy, and yellow). *N* represents the sample size for both groups. Below, the results from a generalized linear mixed model (GLMM) used to test the significance of the small difference in developmental time, with color phenotype as the fixed effect and family as a random effect to control for the genetic similarities between siblings. Since *p* > 0.05, the difference in developmental time is not significant. The data is from a multiyear pedigree of the wood tiger moth laboratory stock maintained by Johanna Mappes’ group. The data and R code for generating both the summary statistics and statistical test can be found in the same Dryad repository as the other materials.

| Color phenotype | N | | Developmental time (days) | | | SE | | SD |
| --- | --- | --- | --- | --- | --- | --- | --- | --- |
| White | 2310 | | 45.178 | | | 0.134 | | 6.441 |
| Yellow | 2039 | | 44.551 | | | 0.137 | | 6.201 |
| **Results from statistical test (GLMM)** | | | | | | | | |
| Fixed effect | | Estimate | | SE | Z | | p-value | |
| Color phenotype | | - 0.002 | | 0.005 | - 0.4 | | 0.666 | |
